# MG53 is not a critical regulator of insulin signaling pathway in skeletal muscle

**DOI:** 10.1101/2020.12.24.424288

**Authors:** Clothilde Philouze, Sophie Turban, Béatrice Cremers, Audrey Caliez, Gwladys Lamarche, Catherine Bernard, Nicolas Provost, Philippe Delerive

**Affiliations:** Institut de Recherches Servier, Cardiovascular and Metabolic Research, 11 rue des Moulineaux, 92150 Suresnes, France

**Keywords:** insulin resistance, ubiquitin E3 ligase, skeletal muscle

## Abstract

In type 2 diabetes (T2D), both muscle and liver are severely resistant to insulin action. Muscle insulin resistance accounts for more than 80% of the impairment in total body glucose disposal in T2D patients and is often characterized by an impaired insulin signaling. Mitsugumin 53 (MG53), a muscle-specific TRIM family protein initially identified as a key regulator of cell membrane repair machinery has been suggested to be a critical regulator of muscle insulin signaling pathway by acting as ubiquitin E3 ligase targeting both the insulin receptor and insulin receptor substrate 1 (IRS1). Here, we show using *in vitro* and *in vivo* approaches that MG53 is not a critical regulator of insulin signaling and glucose homeostasis. First, MG53 expression is not consistently regulated in skeletal muscle from various preclinical models of insulin resistance. Second, MG53 gene knock-down in muscle cells does not lead to impaired insulin response as measured by Akt phosphorylation on Serine 473 and glucose uptake. Third, recombinant human MG53 does not alter insulin response in both differentiated C2C12 and human skeletal muscle cells. Fourth, ectopic expression of MG53 in HEK293 cells lacking endogenous MG53 expression fails to alter insulin response as measured by Akt phosphorylation. Finally, both male and female mg53 −/− mice were not resistant to high fat induced obesity and glucose intolerance compared to wild-type mice. Taken together, these results strongly suggest that MG53 is not a critical regulator of insulin signaling pathway in skeletal muscle.

## Introduction

Type 2 diabetes (T2D) is a global epidemic affecting more than 370 million people worldwide. It is a systemic and progressive disease characterized by hyperglycemia arising, at least in part, from beta cell dysfunction and peripheral insulin resistance [1,2]. Obesity is a major risk factor for T2D [3]. Indeed, more than 80% of patients with T2D are overweight or obese which is a major root cause for the development of insulin resistance [4]. Insulin mediates its physiological effects through the binding to its cognate receptor, namely the insulin receptor (INSR) at the plasma membrane level of target cells (See Petersen and Shulman for review [5]). There are two INSR isoforms, A and B. The B isoform is the primary isoform differentially expressed in insulin responsive tissues such as the liver, adipose tissue and skeletal muscle [6]. In skeletal muscle, insulin binding to INSR triggers the activation of the insulin receptor tyrosine kinase leading to the subsequent phosphorylation of the insulin receptor substrate 1 (IRS-1). Phosphorylated IRS1 recruits and activates phosphatidylinositol-3 kinase leading ultimately via multiple signaling molecules including AKT and RAC1 to the translocation of the glucose transporter type 4 (GLUT-4) to the plasma membrane and the subsequent glucose uptake (See Petersen & Shulman for review [5]). The reduction in membrane INSR levels as well as the impaired downstream signaling contribute to explain the insulin resistance triggered by obesity. Therefore, a better understanding of these complex regulations under pathophysiological conditions may lead to the discovery of novel therapies for the treatment of insulin resistance in T2D.

Two groups reported the identification of Mitsugumin 53 (MG53) as a novel regulator of insulin signaling in skeletal muscle [7,8]. MG53 has been originally identified as a muscle-specific TRIM family protein regulating cell membrane repair machinery [9]. MG53 expression increases during myogenesis and promotes membrane repair by interacting with dysferlin-1 and caveolin-3 [9]. MG53 has emerged as an exciting target for membrane repair. MG53-deficient mice have clear defect in membrane repair in striated muscle resulting in progressive myopathy [9]. Conversely, intravenous injection of recombinant MG53 improves skeletal muscle damage in both mdx [10] and cardiotoxin-injected mice [11]. This therapeutic potential has been later extended to other diseases such as acute kidney injury [12] and acute lung injury [13]. Interestingly, two groups reported that MG53 regulates insulin signaling pathway *in vitro* and more importantly *in vivo* [7,8]. Song and coworkers showed that MG53-deficient mice are resistant to diet-induced obesity and glucose intolerance. Moreover, they showed that mild MG53 over-expression (2 to 3-fold) leads to glucose intolerance and insulin resistance. This was associated to obesity, hypertension and dyslipidemia [8]. Mechanistically, MG53 negatively impacts insulin signaling pathway by targeting both INSR β subunit and IRS-1 protein degradation via its ubiquitin E3 ligase activity [8]. These results were partially confirmed by Yi and colleagues who reported that MG53 induces IRS1 but not INSR b ubiquitination leading to a negative regulation of insulin signaling pathway *in vitro* and *in vivo* [7]. However, MG53 expression was not found altered in preclinical models of insulin resistance and diabetes as well as in patients [7] in contrast to what was reported by Song and colleagues [8]. Nevertheless, results derived from both studies strongly suggested that MG53 might be an interesting target for the treatment off insulin resistance and its associated complications. More recently, Wu and colleagues reported that MG53 is a glucose-sensitive myokine that controls whole body insulin sensitivity by targeting allosterically the insulin receptor [14]. Furthermore, they showed that monoclonal antibody neutralizing circulating MG53 improves hyperglycemia and enhances insulin sensitivity in *db/db* mice [14]. Taken together, these results indicate that MG53 may control insulin sensitivity via multiple mechanisms.

In order to determine the therapeutic potential of targeting MG53 for the treatment of metabolic diseases, we analyzed its regulation across preclinical models of insulin resistance. We next performed both loss and gain of functions in skeletal muscle cells. Since we were not able to confirm the role of MG53 in the regulation of insulin signaling *in vitro*, we engineered mg53 −/− to probe the hypothesis *in vivo*. However, in our hands, mg53 gene deficiency did not protect from diet-induced obesity nor from glucose intolerance. Taken together, our data raise significant doubts about a major role of MG53 in the regulation of insulin signaling pathway and more broadly in glucose homeostasis.

## Materials and Methods

### Reagents

Recombinant human MG53 (GST-tagged) was purchased from Cyclex (Japan, # CY-R2073). The plasmid encoding GFP-tagged TRIM72 (GFP-MG53) and the corresponding empty vector (GFP-Turbo) were purchased from OriGene Technologies, Inc. (Rockville, USA). Insulin was provided by Sigma Aldrich (St QuentinFallavier, France).

### Cell culture

Cell lines were maintained at 37°C and 5% CO2. C2C12 cells (ATCC® CRL-1772™)), a mouse myoblast cell line was cultured in Dulbecco’s Modified Eagle’s Medium supplemented with 10% fetal bovine serum. For differentiation, C2C12 were cultured in the same medium with 2% horse serum for 8 days. Human Skeletal Muscle Myoblasts (HSMM) (CC-2580, Lonza) were cultured following provider’s instructions (Lonza). Finally, HEK293 (ATCC CRL-1573) cells were cultured in Modified Eagle’s Medium (glucose 1G/L) supplemented with 10% fetal bovine serum, sodium pyruvate and non-essential amino acids.

### MG53 gene silencing using RNAi

Twenty-one-nucleotide RNA oligonucleotides directed against mouse MG53 and the non-silencing control siRNA ON-Target plus mouse TRIM72 siRNA-Smart pool L-065686-01 and ON-Target plus Human TRIM72 siRNA-Smart pool L-032293-02 were obtained from Dharmacon™ (Horizon Discovery LTD, Cambridge, UK). C2C12 cells (40% confluence) were transfected with siRNAs (100nM) using Invitrogen™ Lipofectamine™ RNAiMAX transfection reagent (ThermoFisher Scientific, Waltham, MA, USA) following manufacturer’s instructions. 24 hours post-transfection, cells were refed with fresh medium for additional 24 hours.

### MG53 over-expression studies

HEK293 cells, plated in 12-well plates at 50-60% confluence, were transiently transfected with GFP-MG53 or empty vector (GFP-Turbo) (1μg/well). 24h or 48h later, cells were refed with fresh medium containing insulin (100nM) or vehicle (BSA) for 15 minutes. At the end of the treatment period, total cell extracts were prepared for western blot analysis.

### Glucose uptake assay

Differentiated C2C12 cells cultured in 24-well plates were incubated in low glucose medium for 3 hours followed by glucose-free media with or without insulin (100nM) for 30 min. 1 μCi of 2-Deoxy-D-[1-2-^3^H] glucose (PerkinElmer, Boston, MA, USA # NET328A001MC) and 10μM non-radioactive 2-DG (Sigma Aldrich) were added to each well and incubated for 10 min at room temperature. Plates were washed three times in ice-cold PBS and cells were lysed in 500μL of NAOH (1N). ^3^H radioactivity was assessed by scintillation counting on liquid scintillation analyzer Perkin Elmer TriCarb 2910 TR. Total protein levels were quantified to normalize between each well using Bradford reagent (Bio-Rad, Marnes-la-Coquette, France).

### Generation of mg53 −/− mice

The model design and generation have been performed by genOway (Lyon, France). Genomic region of interest containing the murine Trim72 locus was isolated by PCR from C57BL/6 ES cell genomic DNA. PCR fragments were sub-cloned into the pCR4-TOPO vector (Invitrogen, Carlsbad, California). A 4.0-kb region comprising exons 1 and exon 2 (according to ENSEMBL gene structure ENSMUSG00000042828) as well as Trim72 proximal promoter was flanked by a distal loxP site and a Neo cassette (FRT site-PGK promoter-Neo cDNA-FRT site-LoxP site). Transcription and translation start sites as well as the Ring and B-Box Zn finger domains are deleted leading to absence of Trim72 transcription. Linearized targeting vector was transfected into C57BL/6 ES cells (genOway, Lyon, France) according to genOway’s electroporation procedures (ie 108 ES cells in presence of 100μg of linearized plasmid, 260Volt, 500μF). Positive selection was started 48 hours after electroporation, by addition of 200μg/ml of G418 (150μg/ml of active component, Life Technologies, Inc.). G418-Resistant clones were isolated and amplified in 96-well plates. Duplicates of 96-well plates were made. The set of plates containing ES cell clones amplified on gelatin were genotyped by PCR analysis. To fully verify the integrity of the targeted region, the locus, as well as a minimum of 1kb downstream and upstream of both homology arms, was sequenced, confirming the correct targeting event. Clones were microinjected into albino C57BL/6J-Tyrc-2J/J blastocysts, and gave rise to male chimeras with a significant ES cell contribution (as determined by black coat color). Mice were bred to C57BL/6 mice expressing Flpe recombinase to remove the Neo cassette (Trim72lox mice) and to C57BL/6 mice expressing Cre recombinase to generate a constitutive deletion of Trim72 (Trim72del mice). Trim72 gene deletion was verified by PCR and ultimately by western blot analysis. For the sake of clarity, Trim72del mice are referred to mg53 −/− mice throughout the manuscript.

### Animal Studies

Experimental protocols were approved by the Servier Institutional Animal Care and Use Committee. Mice were housed in a specific-pathogen free animal facility with standard 12:12 dark-light cycle with access to standard chow and water ad libitum. Both male and female mg53 −/− (n=6 and 9 per group, respectively) or wild-type mice (n=6 and 9 per group, respectively) were placed under a 60% fat diet (Research Diet # D12492) for 16 weeks. Body weight was recorded on a weekly basis. At the end of the protocol following 4h of fasting, mice were subjected to oral glucose tolerance test (GTT). Briefly, baseline glucose was measured in all animals using a drop of blood from a tail snip wound and Accu-check active glucometers and test strips (Roche Diagnostics). Then, mice received a dose of 2g/kg of D-glucose (Merck) by oral route and blood glucose was monitored at the indicated time points.

Animals were sacrificed the following day and blood was recovered for serum preparation and tissue samples were quickly collected, frozen in liquid nitrogen and used for RNA or total protein extraction.

### RNA analysis

Total RNA was extracted using Qiagen RNA extraction kits following manufacturer’s instructions. Total RNA was treated with DNase I (Ambion Inc., Austin, Texas, USA) at 37°C for 30 minutes, followed by inactivation at 75°C for 5 minutes. Real time quantitative PCR (RT-QPCR) assays were performed using an Applied Biosystems 7500 sequence detector. Total RNA (1 μg) was reverse transcribed with random hexamers using Hight-Capacity cDNA Reverse Transcription Kit with Rnase Inhibitor (Applied Biosystems, ThermoFisher Scientific) following the manufacturer’s protocol. Gene expression levels were determined by Sybr green assays (Mm_Trim72_1_SG QT00315959) using QuantiFast SYBR Green PCR Kit (Qiagen). 18S transcript was used as an internal control to normalize the variations for RNA amounts (Mm_Rn18S_3_SG QT02448075). Gene expression levels are expressed relative to 18S mRNA levels. All the results presented are expressed as mean ± S.E.M. All the primers used in this study and Ct are available upon request.

### Western blot analysis

Protein extracts (30μg) were fractionated on 10% polyacrylamide gel under reducing conditions (sample buffer containing 10 mM dithiothreitol (DTT)) and transferred onto nitrocellulose membranes. Membranes were blocked with 5% milk or BSA 5% in TBS-T 0.1% for 1h. Membranes were then washed three times with TBS-T 0.1% for 10 minutes and incubated overnight at 4°C in blocking buffer containing primary antibodies. Primary antibodies used: phospho-Akt Ser473 (#4060), total Akt (#4691), Hsp90 (# 4877) (Cell Signaling Technology, Danvers, MA), MG53 (Sigma Aldrich, # SAB2108735), Vinculin (Abcam, #ab73412) and Turbo-GFP (OriGene, # TA150041). After incubation with a secondary peroxidase-conjugated antibody, signals were detected by chemiluminescence (GE Healthcare).

### Statistical analysis

Results are shown as means ± S.E.M. Statistical significance was determined using the Student’s *t* test. Differences with p<0.05 were considered to be statistically significant.

## Results & Discussion

Since MG53 has been shown to be a novel regulator of insulin signaling pathway in skeletal muscle, we first determined its expression by qPCR across various mice models of insulin resistance (Fig 1). Interestingly, MG53 was not regulated in a consistent manner across models and muscle types (gastrocnemius versus soleus) (Fig 1 A&B). Its expression is significantly lower in gastrocnemius from both HFD and *ob/ob* mice while there is no change in soleus. By contrast, MG53 was found significantly up-regulated in both muscle types in *db/db* mice. These inconsistent results are in line with previous findings reported by [7,15]. We tried to measure MG53 levels in plasma samples from these animals. However, western blot analyses led to the detection of MG53 at a lower size (data not shown) with doubts about the accuracy of the band for the MG53 signal in line with observations made by other groups [16–18].

**Fig 1.**
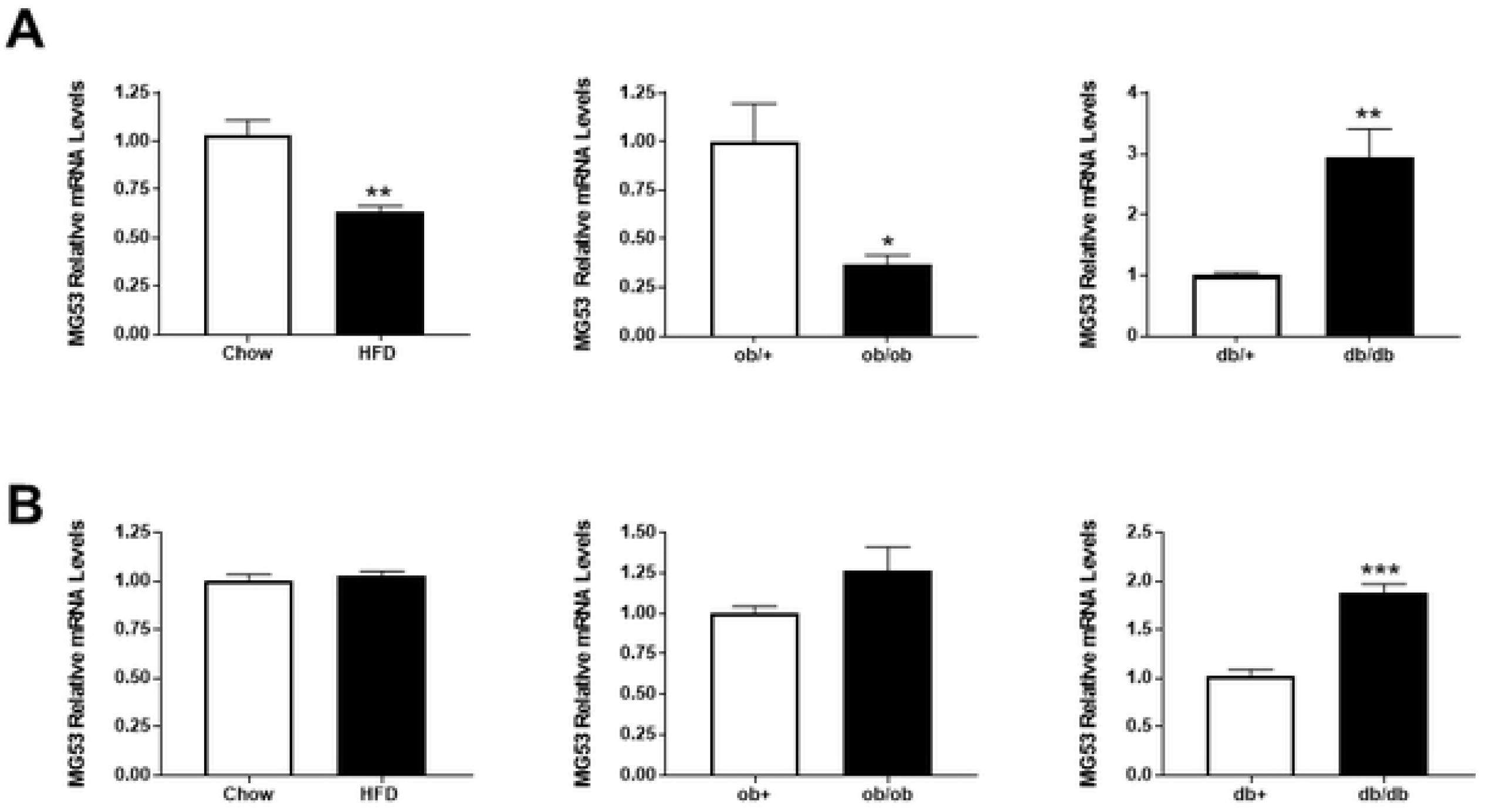
MG53 mRNA levels in skeletal muscles from various preclinical models of insulin resistance. MG53 relative mRNA levels in gastrocnemius (Panel A) or soleus (Panel B) from various preclinical models of insulin resistance: High fat diet (60% fat for 12 weeks), *ob/ob* (10-week old) and *db/db* (10-week old). Data are expressed as means ± SEM (n=10 per group; *: p<0.05, **: p<0.01, ***: p<0.001 diseased *vs.* control).

In order to test directly the role of MG53 in insulin signaling *in vitro*, we performed loss of function experiments using small interfering RNA in differentiated C2C12 cells (Fig 2). Transfection of RNAi targeting MG53 led to a robust and reproducible gene knock-down (−80% p<0.05) compared to siCTL or Mock transfected cells (Fig 2A) which resulted in a complete reduction in MG53 protein levels in C2C12 cells (Fig 2B). In order to evaluate the impact of MG53 gene silencing on insulin signaling pathway, we measured AKT phosphorylation as proxy in response to insulin. As expected, insulin triggered a rapid phosphorylation of AKT on Serine 473 without affecting total AKT levels in siCTL-transfected C2C12 cells (Fig 2C). Interestingly, a similar response to insulin was obtained in siMG53-transfected cells. To assess the functional consequences of insulin signaling activation, we next measured glucose uptake in differentiated cells. Again, insulin triggered as expected a modest but reproducible increase in glucose uptake in mock-transfected (+64%) or siCTL-transfected (+40%) C2C12 cells (Fig 2D). Transfection of siRNA-targeting MG53 did not result in a significant potentiation of insulin response on glucose uptake in differentiated C2C12 cells (+52%) compared to control cells (mock or siCTL-transfected) (Fig 2D). These data are in stark contrast with results reported previously [7,8]. Similar data were obtained in HSMMs using a similar approach. Validated siRNA-targeting MG53 did not alter insulin-response as measured by increased AKT phosphorylation levels (data not shown).

**Fig 2.**
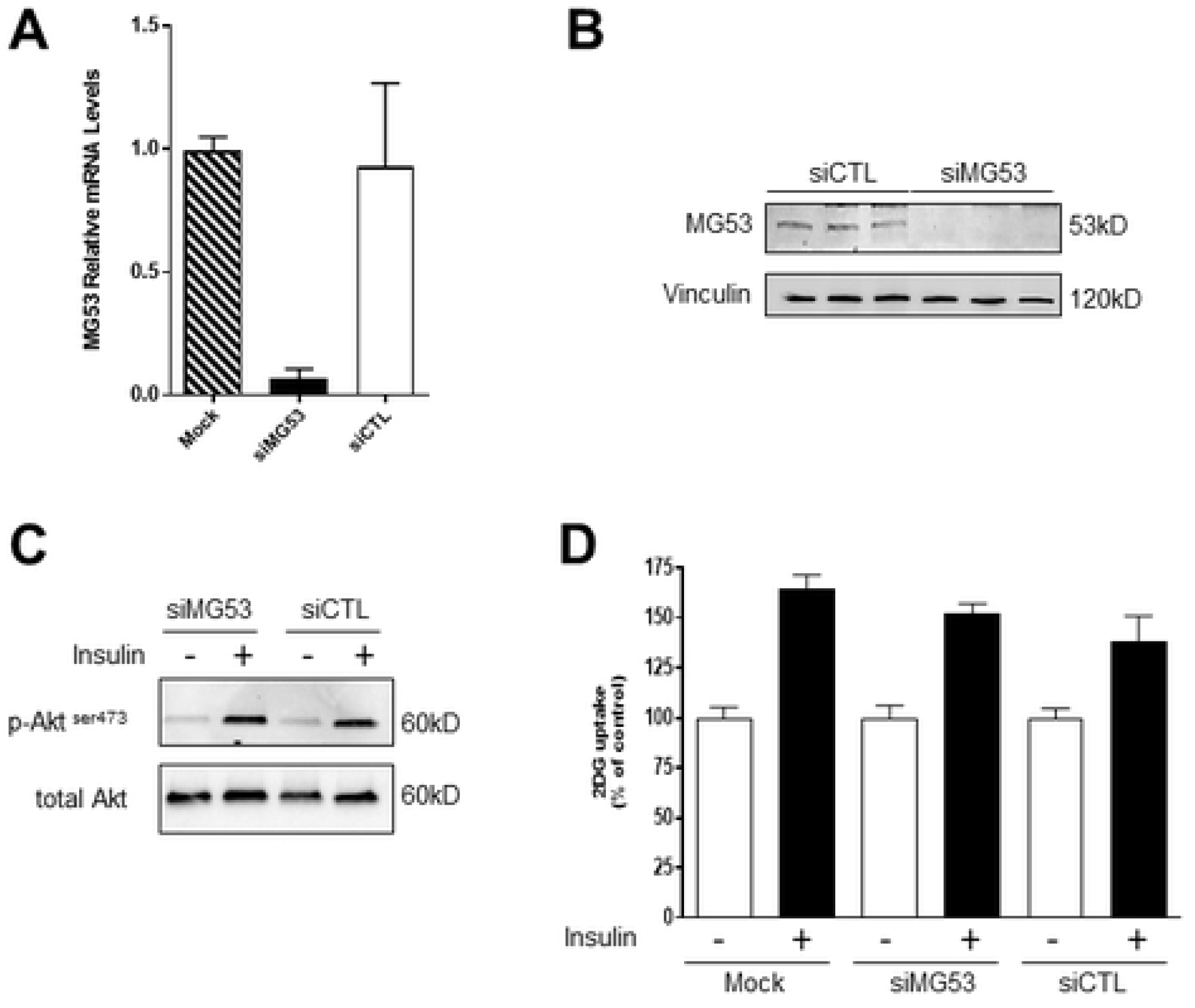
MG53 gene knock-down does not affect insulin signaling in differentiated C2C12 cells. C2C12 cells were transfected with small interfering RNA targeting MG53 or a non-silencing control siRNA (100nM) using Lipofectamine RNAi max. 48h post-transfection, MG53 mRNA (Panel A) and protein levels (Panel B) were assessed by qPCR and western blot analysis, respectively. Vinculin was used as a loading control (Panel B). Insulin response (20nM for 10 minutes) was assessed by monitoring Akt phosphorylation on Serine 473 (Panel C). Finally, glucose uptake was measured using 2DG-glucose in response to insulin treatment (100nM) (Panel C). Data are expressed as percentage of control (Mock-transfected cells).

Since we failed to confirm the role of MG53 as a negative regulator of insulin signaling pathway using loss-function studies *in vitro*, we carried out gain of function studies *in vitro*. HEK293 cells were selected as a simple cellular system with good response to insulin and devoid of endogenous MG53 expression. MG53 was successfully over-expressed in HEK293 cells by transient transfection using a plasmid encoding a GFP-tagged human MG53 as demonstrated by significant protein levels measured by western blot analysis. This led to a robust MG53 expression 24 and 48h post transfection. As expected, insulin treatment irrespectively of the tested concentrations (10 or 100nM) led to AKT phosphorylation without affecting total AKT levels on both mock and GFP-transfected cells (Fig 3). MG53 over-expression failed to alter this response at both 24h and 48h time points irrespectively of the tested insulin concentrations. Similar results were obtained in HepG2 cells, a human hepatoma cell line lacking also MG53 endogenous expression (data not shown).

**Fig 3.**
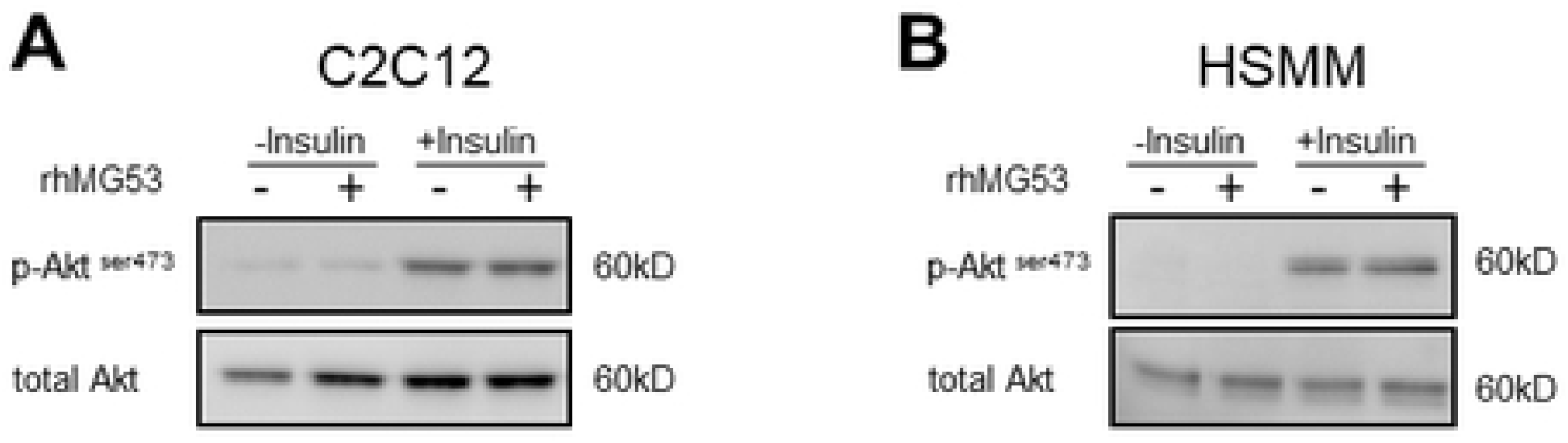
MG53 over-expression does not affect insulin response in HEK293 cells. HEK293 cells were transfected with a MG53 expression plasmid or GFP as control. 24h (left panel) or 48h later (right panel), cells are exposed to increasing concentrations of insulin (10 and 100nM) for 15 minutes. Insulin response was then evaluated using phosphor Akt levels.

Since MG53 has recently been shown to behave as a glucose-responsive myokine able to control peripheral insulin sensitivity by allosterically regulating the insulin receptor [14], we tested human recombinant MG53 protein on insulin-mediated AKT phosphorylation in both mouse (C2C12) and human (HSMMs) skeletal muscle cells (Fig 4). Treatment with recombinant human MG53 (30μg/mL) did not result in a significant inhibition of insulin-mediated AKT phosphorylation in both C2C12 cells and HSMMs (Fig 4 A&B). Concentration-response studies (from 0.1 up to 30μg/mL) and extensive kinetic studies were performed (data not shown). Despite all the conditions tested, these experiments failed to reveal a negative role of MG53 on insulin signaling.

**Fig 4.**
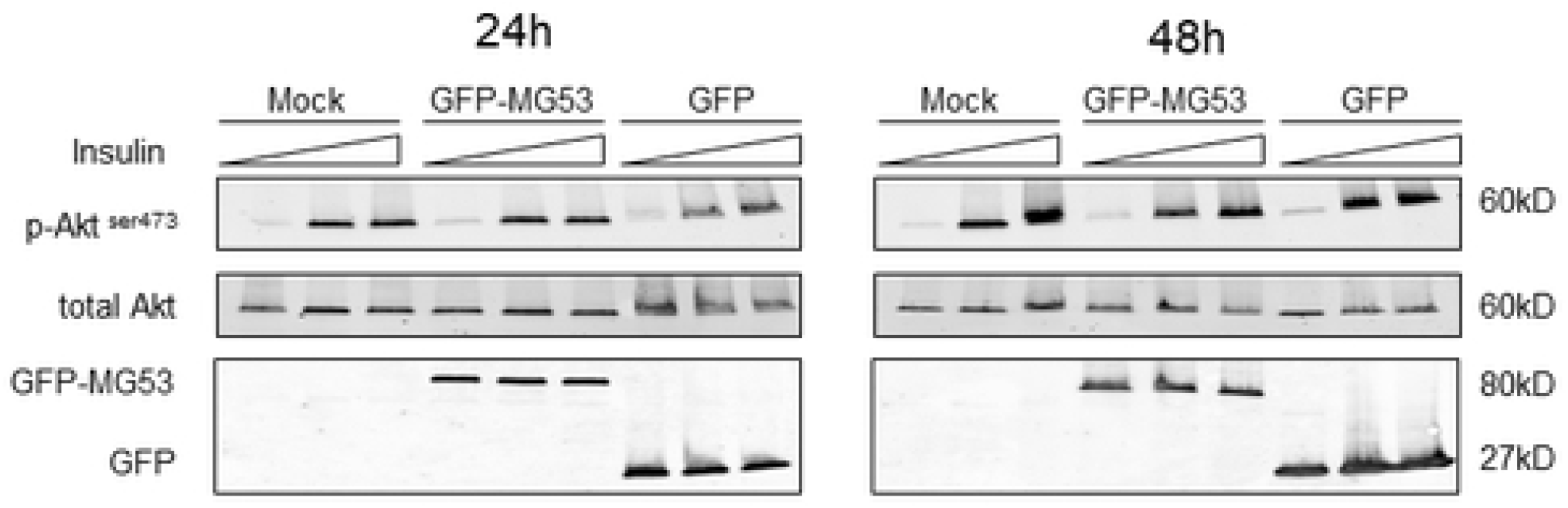
Recombinant human MG53 does not affect insulin response in both C2C12 cells and primary human skeletal muscle myoblasts. Differentiated C2C12 cells (Panel A) or HSMMs (Panel B) were preincubated with rhMG53 (30μg/mL) for 1h before being treated with insulin (20nM for 10 minutes and 100nM for 30 minutes, respectively) or vehicle (BSA). Akt phosphorylation was then evaluated by western blot analysis.

Since all *in vitro* approaches failed to confirm the role of MG53 as a modulator of insulin signaling pathway, we engineered mg53 −/− mice by targeting both exon 1 and 2 (Fig 5A). Both transcription and translation start sites as well as the Ring and B-Box Zn finger domains were deleted leading to absence of mg53 transcription. This approach led to the generation of mg53 −/− mice in C5BL/6J genetic background. These mice did not display any gross abnormalities compared to wild type mice even though mg53 −/− mice were born at a less than expected ratio. In order to directly test the impact of MG53 gene deficiency on glucose homeostasis, both male and female mice (mg53 −/− and wild type) were placed on high fat diet for 16 weeks. Western blot analyses confirmed at the protein level the efficient deletion of MG53 gene in all animals included in the study (Fig 5B). MG53 gene deficiency did not affect weight gain nor total body weight in both males and females compared to wild type mice (Fig 5C). The high fat diet triggered a significant glucose intolerance as revealed by the oral GTT and an elevated fasting glycemia in both male and female wild type mice (Fig 5D) with males being more glucose intolerant as expected. Fasting glycemia and response to oral GTT were similar in both KO and control mice irrespectively of the gender (Fig 5D). These results suggest that MG53 is a not a critical regulator of glucose homeostasis in mice.

**Fig 5.**
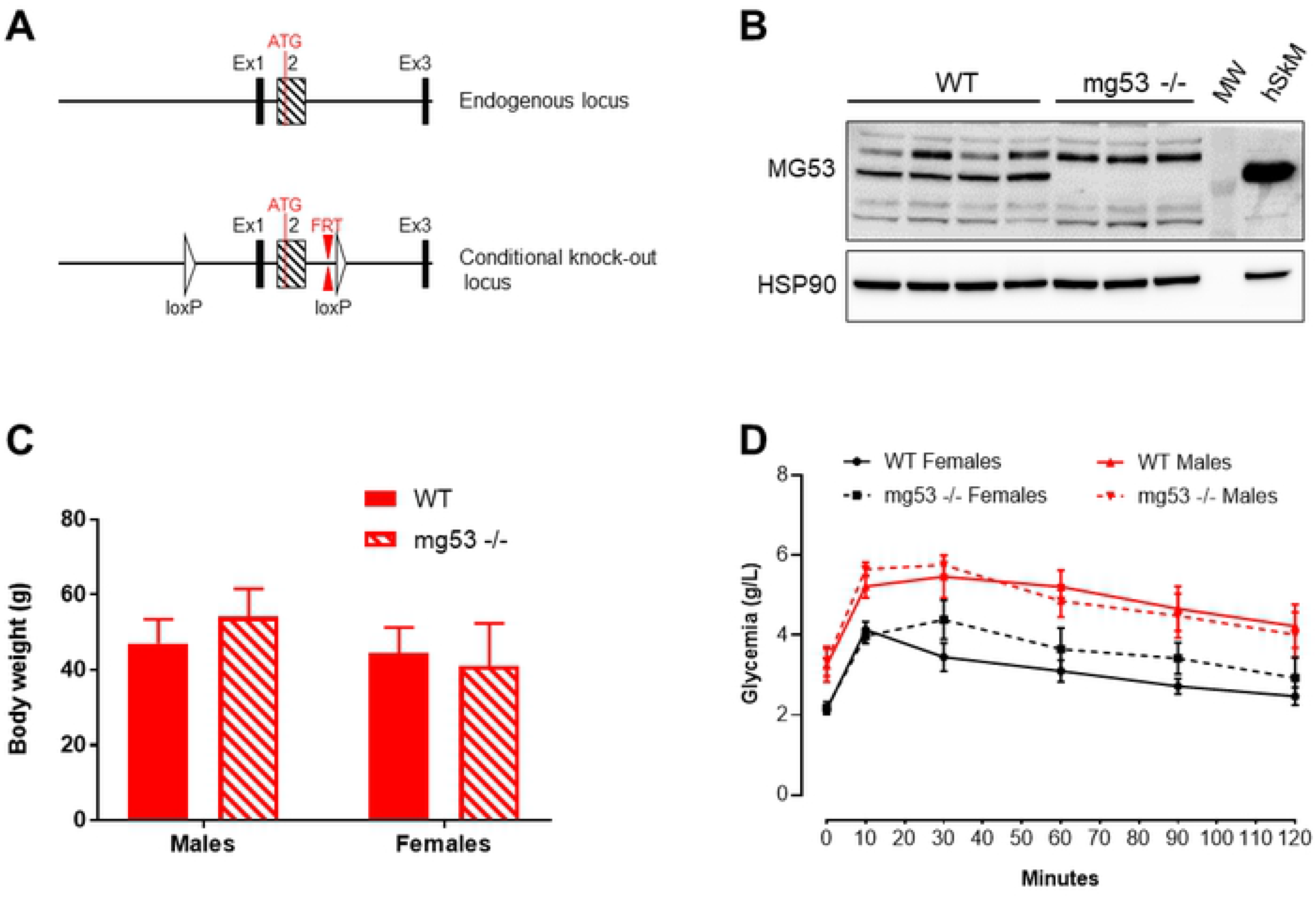
mg53 gene deficiency does not affect high fat diet-induced obesity and glucose intolerance. Panel A: targeting strategy to engineer mg53 −/− mice. Panel B: western blot analysis of MG53 expression in both wild type and KO mice. MW: molecular weight marker. 10μg of total protein lysate from human skeletal muscle was used as positive control. Panel C: Total body weight in both males and females wild-type and mg53 KO mice after 16 weeks of high fat diet. Panel D: An oral GTT was performed in both male and female mice (wild-type vs. KO mice) after 16 weeks of high fat diet.

Our results failed to confirm the role of MG53 as a critical regulator of insulin signaling pathway *in vitro* and of glucose homeostasis *in vivo* in contrast to earlier reports [7,8]. While we cannot exclude the possibility that different reagents may lead to different results, our data raise significant doubts about the involvement of MG53 in the regulation of insulin signaling. In addition, our data are consistent with recent reports from the Ma’s lab who used alternative strategies to come to the same conclusions [16,19]. First, they showed using transgenic mice that sustained elevation of plasma MG53 levels does not compromise insulin signaling in skeletal muscle as well as glucose handling when mice were placed under high fat diet. Furthermore, they reported no impact of sustained elevation of plasma MG53 on the diabetic phenotype in *db/db* mice [19]. They further extended their findings more recently by crossing mg53 −/− with *db/db* mice. MG53 gene deficiency has no impact on total body weight nor on glucose homeostasis in this genetic background [16]. Our *in vitro* and *in vivo* results are highly consistent with their conclusions. In conclusion, these results do not support to further investigate MG53 as a target for insulin resistance and T2D.

The identification of novel regulators of insulin signaling pathway remains of utmost importance for the treatment of T2D and its associated complications. Several E3 ubiquitin ligases have been identified as regulators of insulin signaling pathway [20,21]. More recently, Nagarajan and coworkers identified using a large-scale RNAi screen performed in HeLa cells MARCH1 as a novel negative regulator of insulin signaling [22]. Interestingly, MARCH1 controls this pathway by regulating basal insulin receptor levels via direct ubiquitination. Furthermore, its expression is altered in white adipose tissue from obese humans suggesting its potential involvement in the development of insulin resistance [22]. Whether MARCH1 could be an interesting drug target for metabolic diseases remains to be addressed since MARCH1 is known to play a role in airway allergic immunity by, at least in part, mediating ubiquitination of MHCII and CD86 in dendritic cells [23]. Furthermore, its role in CD8 T cell in adipose tissue inflammation and the link to obesity-induced insulin resistance in mice requires further studies to better understand the overall impact of MARCH1 on glucose and energy homeostasis [24].

## Conclusions

Our results derived from *in vitro* and *in vivo* studies raise significant doubts about the role of MG53 in the regulation of insulin signaling pathway and more broadly in glucose homeostasis.

## Acknowledgments

The authors would like to thank Kader Thiam and Marie Norbert from Genoway (Lyon, France) for mg53 −/− mice engineering and Emmanuelle Douillet, Mathilde Lecomte, Christophe Ragonnet and Kevin Romariz for excellent technical assistance.

## References

1. DeFronzo RA, Tripathy D. Skeletal muscle insulin resistance is the primary defect in type 2 diabetes. Diabetes care. Diabetes Care; 2009. doi:10.2337/dc09-s302

2. Defronzo RA. From the triumvirate to the ominous octet: A new paradigm for the treatment of type 2 diabetes mellitus. Diabetes. Diabetes; 2009. pp. 773–795. doi:10.2337/db09-9028

3. Seshasai SRK, Kaptoge S, Thompson A, Di Angelantonio E, Gao P, Sarwar N, et al. Diabetes mellitus, fasting glucose, and risk of cause-specific death. N Engl J Med. 2011;364: 829–841. doi:10.1056/NEJMoa1008862

4. Shulman GI. Ectopic fat in insulin resistance, dyslipidemia, and cardiometabolic disease. New England Journal of Medicine. Massachussetts Medical Society; 2014. pp. 1131–1141. doi:10.1056/NEJMra1011035

5. Petersen MC, Shulman GI. Mechanisms of insulin action and insulin resistance. Physiological Reviews. American Physiological Society; 2018. pp. 2133–2223. doi:10.1152/physrev.00063.2017

6. Belfiore A, Frasca F, Pandini G, Sciacca L, Vigneri R. Insulin receptor isoforms and insulin receptor/insulin-like growth factor receptor hybrids in physiology and disease. Endocrine Reviews. Endocr Rev; 2009. pp. 586–623. doi:10.1210/er.2008-0047

7. Yi JS, Park JS, Ham YM, Nguyen N, Lee NR, Hong J, et al. MG53-induced IRS-1 ubiquitination negatively regulates skeletal myogenesis and insulin signalling. Nat Commun. 2013;4: 1–12. doi:10.1038/ncomms3354

8. Song R, Peng W, Zhang Y, Lv F, Wu HK, Guo J, et al. Central role of E3 ubiquitin ligase MG53 in insulin resistance and metabolic disorders. Nature. 2013;494: 375–379. doi:10.1038/nature11834

9. Cai C, Masumiya H, Weisleder N, Matsuda N, Nishi M, Hwang M, et al. MG53 nucleates assembly of cell membrane repair machinery. Nat Cell Biol. 2009;11: 56–64. doi:10.1038/ncb1812

10. Weisleder N, Takizawa N, Lin P, Wang X, Cao C, Zhang Y, et al. Recombinant MG53 protein modulates therapeutic cell membrane repair in treatment of muscular dystrophy. Sci Transl Med. 2012;4: 139ra85–139ra85. doi:10.1126/scitranslmed.3003921

11. Zhang Y, Wu HK, Lv F, Xiao RP. MG53: Biological function and potential as a therapeutic target. Molecular Pharmacology. American Society for Pharmacology and Experimental Therapy; 2017. pp. 211–218. doi:10.1124/mol.117.108241

12. Duann P, Li H, Lin P, Tan T, Wang Z, Chen K, et al. MG53-mediated cell membrane repair protects against acute kidney injury. Sci Transl Med. 2015;7: 279ra36–279ra36. doi:10.1126/scitranslmed.3010755

13. Jia Y, Chen K, Lin P, Lieber G, Nishi M, Yan R, et al. Treatment of acute lung injury by targeting MG53-mediated cell membrane repair. Nat Commun. 2014;5: 1–12. doi:10.1038/ncomms5387

14. Wu H-K, Zhang Y, Cao C-M, Hu X, Fang M, Yao Y, et al. Glucose-Sensitive Myokine/Cardiokine MG53 Regulates Systemic Insulin Response and Metabolic Homeostasis. Circulation. 2019;139: 901–914. doi:10.1161/CIRCULATIONAHA.118.037216

15. Ma H, Liu J, Bian Z, Cui Y, Zhou X, Zhou X, et al. Effect of Metabolic Syndrome on Mitsugumin 53 Expression and Function. Peterson J, editor. PLoS One. 2015;10: e0124128. doi:10.1371/journal.pone.0124128

16. Wang Q, Bian Z, Jiang Q, Wang X, Zhou X, Park KH, et al. Mg53 does not manifest the development of diabetes in db/db mice. Diabetes. 2020;69: 1052–1064. doi:10.2337/db19-0807

17. Zhu H, Hsueh W, Whitson BA. Letter by Zhu et al Regarding Article, “glucose-Sensitive Myokine/Cardiokine MG53 Regulates Systemic Insulin Response and Metabolic Homeostasis.” Circulation. Lippincott Williams and Wilkins; 2019. pp. E186–E187. doi:10.1161/CIRCULATIONAHA.118.039305

18. Lemckert FA, Bournazos A, Eckert DM, Kenzler M, Hawkes JM, Butler TL, et al. Lack of MG53 in human heart precludes utility as a biomarker of myocardial injury or endogenous cardioprotective factor. Cardiovasc Res. 2016;110: 178–187. doi:10.1093/cvr/cvw017

19. Bian Z, Wang Q, Zhou X, Tan T, Park KH, Kramer HF, et al. Sustained elevation of MG53 in the bloodstream increases tissue regenerative capacity without compromising metabolic function. Nat Commun. 2019;10: 1–16. doi:10.1038/s41467-019-12483-0

20. Rui L, Yuan M, Frantz D, Shoelson S, White MF. SOCS-1 and SOCS-3 block insulin signaling by ubiquitin-mediated degradation of IRS1 and IRS2. J Biol Chem. 2002;277: 42394–42398. doi:10.1074/jbc.C200444200

21. Xu X, Sarikas A, Dias-Santagata DC, Dolios G, Lafontant PJ, Tsai SC, et al. The CUL7 E3 Ubiquitin Ligase Targets Insulin Receptor Substrate 1 for Ubiquitin-Dependent Degradation. Mol Cell. 2008;30: 403–414. doi:10.1016/j.molcel.2008.03.009

22. Nagarajan A, Petersen MC, Nasiri AR, Butrico G, Fung A, Ruan H Bin, et al. MARCH1 regulates insulin sensitivity by controlling cell surface insulin receptor levels. Nat Commun. 2016;7: 1–16. doi:10.1038/ncomms12639

23. Corcoran K, Jabbour M, Bhagwandin C, Deymier MJ, Theisen DL, Lybarger L. Ubiquitin-mediated regulation of CD86 protein expression by the ubiquitin ligase membrane-associated RING-CH-1 (MARCH1). J Biol Chem. 2011;286: 37168–37180. doi:10.1074/jbc.M110.204040

24. Majdoubi A, Lee JS, Kishta OA, Balood M, Moulefera MA, Ishido S, et al. Lack of the E3 Ubiquitin Ligase March1 Affects CD8 T Cell Fate and Exacerbates Insulin Resistance in Obese Mice. Front Immunol. 2020;11: 1953. doi:10.3389/fimmu.2020.01953

